# 1,25(OH)_2_D_3_ attenuates hepatic insulin resistance in rats with streptozotocin-induced diabetes via antioxidation

**DOI:** 10.1101/2021.07.27.453999

**Authors:** Guoyu Huang, Runrong Ding, Yujing Zhang, Jiaojiao Gao, Ze Xu, Wenjie Li, Xing Li

## Abstract

**Aim:** This study was conducted to explore the mechanism by which 1,25(OH)_2_D_3_ ameliorates hepatic insulin resistance in rats with streptozotocin (STZ)-induced type 2 diabetes.

**Methods:** Sprague Dawley (SD) rats were randomly divided into five groups: normal, control, and 1,25(OH)_2_D_3_ administered at 0.075, 0.15, and 0.3 μg/kg/d. Tissue and blood samples were obtained from all five groups after 4 and 12 weeks of treatment. Morphological changes in the livers were observed with HE staining. The levels of fasting blood glucose (FBG), insulin, and lipids were determined using an automatic biochemistry analyzer. The levels of hepatic glycogen, superoxide dismutase (SOD), malondialdehyde (MDA), aspartate aminotransferase (AST), and alanine aminotransferase (ALT) were analyzed by spectrophotometry. Protein expression of AKT and GLUT2 was examined by western blotting.

**Results:** 1,25(OH)_2_D_3_ treatment reduced the FBG content as well as hepatic MDA, AST, and ALT levels, improved the activity of SOD, and enhanced the expression of GLUT2 and AKT in diabetic rats. These findings suggest that 1,25(OH)_2_D_3_, exerts antioxidant effects and ameliorates insulin resistance in the liver.

**Conclusion:** 1,25(OH)_2_D_3_ is a potential agent that could be used in the treatment of insulin resistance in the liver.

## Introduction

Diabetes mellitus (DM) is a group of chronic, systemic, and metabolic diseases characterized by hyperglycemia, which is caused by insufficient insulin secretion and/or insulin resistance (IR) occurring due to genetic and environmental factors [1]. Oxidative stress has long been considered one of the main damaging factors responsible for the development of insulin resistance, impaired insulin secretion, and pathogenesis of type 2 diabetes mellitus (T2DM) [2]. The liver is an important organ where the synthesis and metabolism of carbohydrates, fats, and proteins occur; these processes are regulated by insulin under normal physiological conditions [3]. Insulin resistance in the liver increases the risk of disorders of glucose-lipid metabolism [4].

Vitamin D is a fat-soluble vitamin necessary for maintaining human life. Vitamin D plays a role in calcium-phosphorus regulation and bone metabolism; moreover, abnormal levels of vitamin D have been linked to several other diseases, such as cancer and chronic diseases, including obesity, cardiovascular disease, diabetes, and insulin resistance-related diseases [5,6,7,8]. Insulin resistance is known to be associated with vitamin D deficiency, and supplementation with 1,25(OH)_2_D_3_ is known to improve insulin resistance; however, the underlying mechanism remains unclear. In the current study, using type 2 diabetic rats, we explored the mechanistic role of 1,25(OH)_2_D_3_ in ameliorating hepatic insulin resistance.

## Materials and methods

### Reagents

Streptozotocin (STZ) was obtained from Sigma-Aldrich (St. Louis, MO, USA). 1,25(OH)_2_D_3_ was purchased from Roche Pharma (Basel, Switzerland). Insulin, total cholesterol (TC), triglyceride (TG), high density lipoprotern-cholesterol (HDL-C) and low density lipoprotern-cholesterol (LDL-C) assay kits were purchased from Zhongsheng Beikong Biological Technology Co. Ltd., China. Superoxide dismutase (SOD) and malondialdehyde (MDA) assay kits were purchased from Jiancheng Institute of Biological Engineering, Nanjing, China. Aspartate aminotransferase (AST) and alanine aminotransferase (ALT) assay kits were purchased from Zhongsheng Beikong Biological Technology Co. Ltd., China. Antibodies for the detection of AKT, GLUT2 and β-actin were purchased from ProteinTech Group, China.

### Animal Models

Sprague Dawley (SD) rats (male, 150 ±20 g, 4-week-old) were purchased from the Laboratory Animal Center of Henan Province. All rats were maintained in 50–60% humidity, a temperature of 23-26 °C, and a 12 h light/dark cycle. All animal experiments were performed in accordance with the National Research Council’s Laboratory Animal Care and Use guidelines. Ten rats were randomly divided into a normal group, which was fed an ordinary diet. The remaining rats were fed a high-fat and high-sugar diet for 5 weeks and then injected with 1% STZ (35 mg/kg) to establish the diabetes model. The diabetic rats were randomly divided into four groups: control group (diabetic rats, n=12), low-dose group (gavage of 0.075 μg/kg/d 1,25(OH)_2_D_3_; n=12), middle-dose group (gavage of 0.150 μg/kg/d 1,25(OH)_2_D_3_; n=12), and high-dose group (gavage of 0.3 μg/kg/d 1,25(OH)_2_D_3_; n=12). The intervention duration was divided into 4 and 12 weeks.

### Experimental Analysis

Blood samples were collected, and plasma was separated by centrifugation. After sacrificing the animals, the rat liver was quickly extracted and washed with PBS. Blood and tissue samples were used to analyze glycolipid metabolism, antioxidant stress, liver function enzymes, protein expression, and histopathology.

### Estimation glucose and lipid metabolism indexes

Detection of insulin, TG, TC, HDL-C, LDL-C and FBG in serum by automatic biochemical analyzerLiver glycogen was evaluated by colorimetry.

### Detection of biological indicators of oxidative stress

The contents of SOD and MDA in rat liver were determined by colorimetry according to the operation method of the kit.

### Liver function indicator

The hepatic marker enzymes like aspartate aminotransferase (AST),alanine aminotransferase (ALT) inserum were estimated using Colorimetric determination.

### Western blotting analysis

The extracted liver tissue extract protein (30 μg) was separated on a 10% SDS-PAGE gel and transferred to a polyvinylidene difluoride (PVDF) membrane. The blotted membrane was blocked with 5% skimmed milk in TBST (Tris Buffered Saline with 10% of tween) buffer for 1.5 hours. The blotted membrane was then incubated with primary antibodies against AKT and GLUT2 were diluted to 1:5000 and 1:500 respectively, at 4 °C overnight, washed thrice with TBST buffer for 10 min and then incubated with a 1:1000 dilution of horseradish peroxidase-conjugated secondary antibody for 120 min. The immunoreactive bands were visualized using a chemiluminescence imager (Nikon Digital Sight DS-FI2). Soft ImageJ (NIH, Maryland, USA) was used to calculate the density values of the bands.

### Histopathology of liver

The liver tissue was fixed in 10% formalin for 48 hours, dehydrated, and then gradually graded from 50% ethanol to 100% (absolute) ethanol through a series of graded alcohols, and finally embedded in paraffin. Semi-automatic rotary microtome scans liver slices (5-6 m thick), stained with hematoxylin and eosin (HE), and observed under microscope [9].

### Statistical Analysis

All data were expressed as means ± SD. Results were analysed by one-way ANOVA and t-test. P < 0.05 was considered significantly different.

## Results

### Changes in body weight, fasting blood glucose (FBG), insulin and liver glycogen levels in different groups

As shown in Table 1, the average body weight of control rats was significantly lower than that of the normal rats. After 4 and 12 weeks of intervention, the body weights of the high-dose groups increased significantly compared to those of the control groups. Compared to the control group, at 4 weeks of intervention, the level of FBG in the low-dose group was decreased, the insulin levels of both low-dose and high-dose groups were significantly increased, and the content of hepatic glycogen in the middle-dose group was significantly increased. Compared to the control group at 12 weeks of intervention, all treatment groups showed reduction in the FBG levels and improvement in the insulin levels. However, no significant difference in liver glycogen levels was observed between the groups.

**Table 1.** Changes in body weight, fasting blood glucose, insulin, and liver glycogen levels in rats after treatment.

|  | Normal | Control | L | M | H |
| --- | --- | --- | --- | --- | --- |
| Body weight(g) |  |  |  |  |  |
| week 0 | 411.47±44.98 | 371.67±44.21 | 372.53±44.21 | 361.63±25.58 | 363.91±39.11* |
|  |  | * | * | * |  |
| week 4 | 435.50±71.64 | 345.27±56.23 | 358.26±50.16 | 367.63±39.80 | 375.65±55.00* |

|  |  | * | * | * | # |
| --- | --- | --- | --- | --- | --- |
| week 12 | 515.63±56.15 | 366.54±65.25 | 366.50±66.51 | 406.54±51.82 | 448.25±92.74<sup>#</sup> |
|  | # | * | * | * |  |
| FBG |  |  |  |  |  |
| week 0 | 5.64±1.05 | 21.37±4.64* | 21.88±5.71* | 21.00±4.68* | 21.73±5.40* |
| week 4 | 6.33±0.45<sup>#</sup> | 27.32±2.86* | 12.21±5.25*<sup>#</sup> | 22.04±8.14* | 26.78±6.11* |
| week 12 | 5.77±0.63<sup>#</sup> | 27.79±2.12* | 19.65±7.07*<sup>#</sup> | 21.86±5.92*<sup>#</sup> | 21.13±4.78*<sup>#</sup> |
| Insulin |  |  |  |  |  |
| week 0 | 3.57±1.68 | 5.43±1.51 | 5.51±2.04 | 5.28±1.07 | 4.95±0.90 |
| week 4 | 2.52±0.76 | 2.54±0.95 | 4.56±0.81*<sup>#</sup> | 3.76±1.13 | 4.00±0.65*<sup>#</sup> |
| week 12 | 3.58±1.24 | 2.29±0.92* | 4.25±0.90<sup>#</sup> | 3.52±0.85<sup>#</sup> | 3.98±1.30<sup>#</sup> |
| Liver glycogen |  |  |  |  |  |
| week 4 | 3.68±2.29<sup>#</sup> | 5.95±0.85* | 4.23±0.52<sup>#</sup> | 4.18±0.68<sup>#</sup> | 6.63±0.89* |
| week 12 | 5.16±1.11 | 4.11±0.50 | 5.64±1.68 | 4.25±0.63 | 5.18±2.13 |

Normal, normal rats; control, type 2 diabetes rats; L, type 2 diabetes rats treated with 0.075 µg/Kg/d of 1,25(OH)<sub>2</sub>D<sub>3</sub>; M, type 2 diabetes rats treated with 0.150 µg/Kg/d of 1,25(OH)<sub>2</sub>D<sub>3</sub>; H, type 2 diabetes rats treated with 0.300 µg/Kg/d of 1,25(OH)<sub>2</sub>D<sub>3</sub>.
\* $p < 0.05$ as compared to normal; # $p < 0.05$ as compared to control

### Changes in TC, TG, HDL-C, and LDL-C in rats after different doses and durations of 1,25(OH)_2_D_3_ treatment

As depicted in Table 2, 4 weeks of 1,25(OH)_2_D_3_ intervention had no obvious effect on serum TG, TC, and LDL-C levels (*P*>0.05). After 12 weeks of intervention, compared to the control group, the serum levels of TC, TG, and LDL-C in the high-dose 1,25(OH)_2_D_3_ group significantly decreased, while the HDL-C levels increased (*P*<0.05).

**Table 2.** Changes in TC, TG, HDL-C and LDL-C in Rats after Different Treatment.

|  | Normal | Control | L | M | H |
| --- | --- | --- | --- | --- | --- |
| TC |  |  |  |  |  |
| week 0 | 1.67±0.15 | 2.15±0.42 | 2.23±0.83 | 2.54±0.56 | 2.27±0.78 |
| week 4 | 1.38±0.12 | 2.32±0.70 | 2.54±1.87 | 2.53±1.03 | 2.57±1.40 |
| week 12 | 1.53±0.20<sup>#</sup> | 2.74±0.48* | 2.18±0.78*<sup>#</sup> | 2.40±0.68* | 1.49±0.38<sup>#</sup> |
| TG |  |  |  |  |  |
| week 0 | 0.85±0.30 | 2.04±1.32* | 3.29±1.63*<sup>#</sup> | 2.45±1.62* | 1.98±1.14* |
| week 4 | 0.63±0.27 | 3.50±1.78* | 1.69±2.03 | 3.85±2.91* | 4.10±3.44* |
| week 12 | 0.77±0.34 | 4.11±1.72* | 2.80±2.30* | 2.99±1.72* | 2.05±2.05<sup>#</sup> |
| HDL-C |  |  |  |  |  |
| week 0 | 1.00±0.11 | 0.63±0.35 | 0.68±0.39 | 0.68±0.33 | 0.63±0.35 |
| week 4 | 1.33±0.52<sup>#</sup> | 0.50±0.25* | 1.11±0.39<sup>#</sup> | 0.95±0.48<sup>#</sup> | 0.91±0.29*<sup>#</sup> |
| 7week 12 | 1.21±0.49 | 0.73±0.26* | 0.83±0.32* | 0.79±0.13* | 1.13±0.19<sup>#</sup> |
| LDL-C |  |  |  |  |  |
| week 0 | 0.32±0.05 | 0.44±0.15 | 0.71±0.67* | 0.59±0.43 | 0.55±0.54 |
| week 4 | 0.36±0.09 | 0.71±0.31 | 1.00±0.59 | 1.31±0.62 | 1.11±1.03 |
| week 12 | 0.54±0.34 | 0.93±0.48 | 0.67±0.53 | 1.14±0.78* | 0.36±0.16<sup>#</sup> |

Normal, normal rats; control, type 2 diabetes rats; L, type 2 diabetes rats treated with 0.075 µg/Kg/d of 1,25(OH)<sub>2</sub>D<sub>3</sub>; M, type 2 diabetes rats treated with 0.150 µg/Kg/d of 1,25(OH)<sub>2</sub>D<sub>3</sub>; H, type 2 diabetes rats treated with 0.300 µg/Kg/d of 1,25(OH)<sub>2</sub>D<sub>3</sub>.
\* $p < 0.05$ as compared to normal; # $p < 0.05$ as compared to control

### Changes in hepatic MDA and SOD levels in rats from different groups

To explore the changes in oxidative stress levels, the levels of hepatic SOD and MDA in all groups were tested (Fig 1). After 4 weeks of intervention, the SOD activity of the middle-dose intervention group was significantly increased, and the MDA levels of the low-dose and high-dose intervention groups were decreased as compared to those of the control group. After 12 weeks of intervention, the SOD activity of all treatment groups was significantly increased, and the MDA levels of the middle-dose and high-dose intervention groups were significantly reduced as compared to those of the control group.

**Fig 1.**
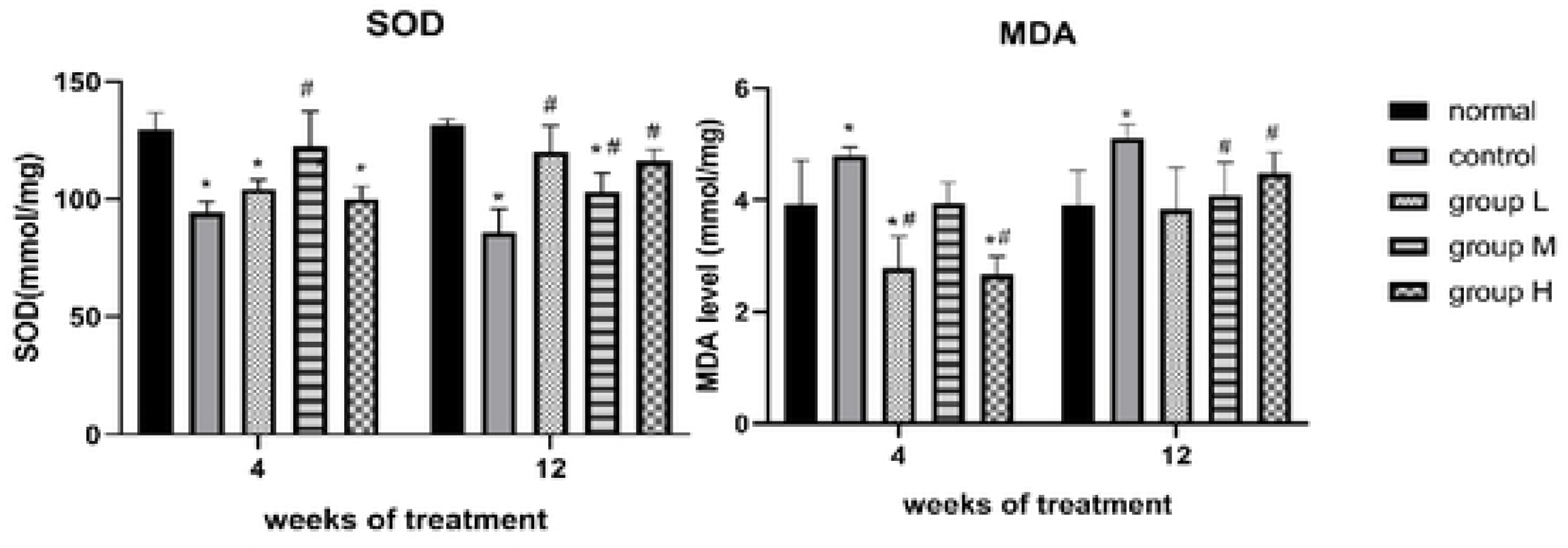

### Changes in ALT, AST in liver

1,25(OH)_2_D_3_ intervention had no obvious effect on the hepatic ALT level of diabetic rats, whereas short-term 1,25(OH)_2_D_3_ intervention was able to reduce the hepatic AST levels of diabetic rats, regardless of the concentrations (Fig 2).

**Fig 2.**
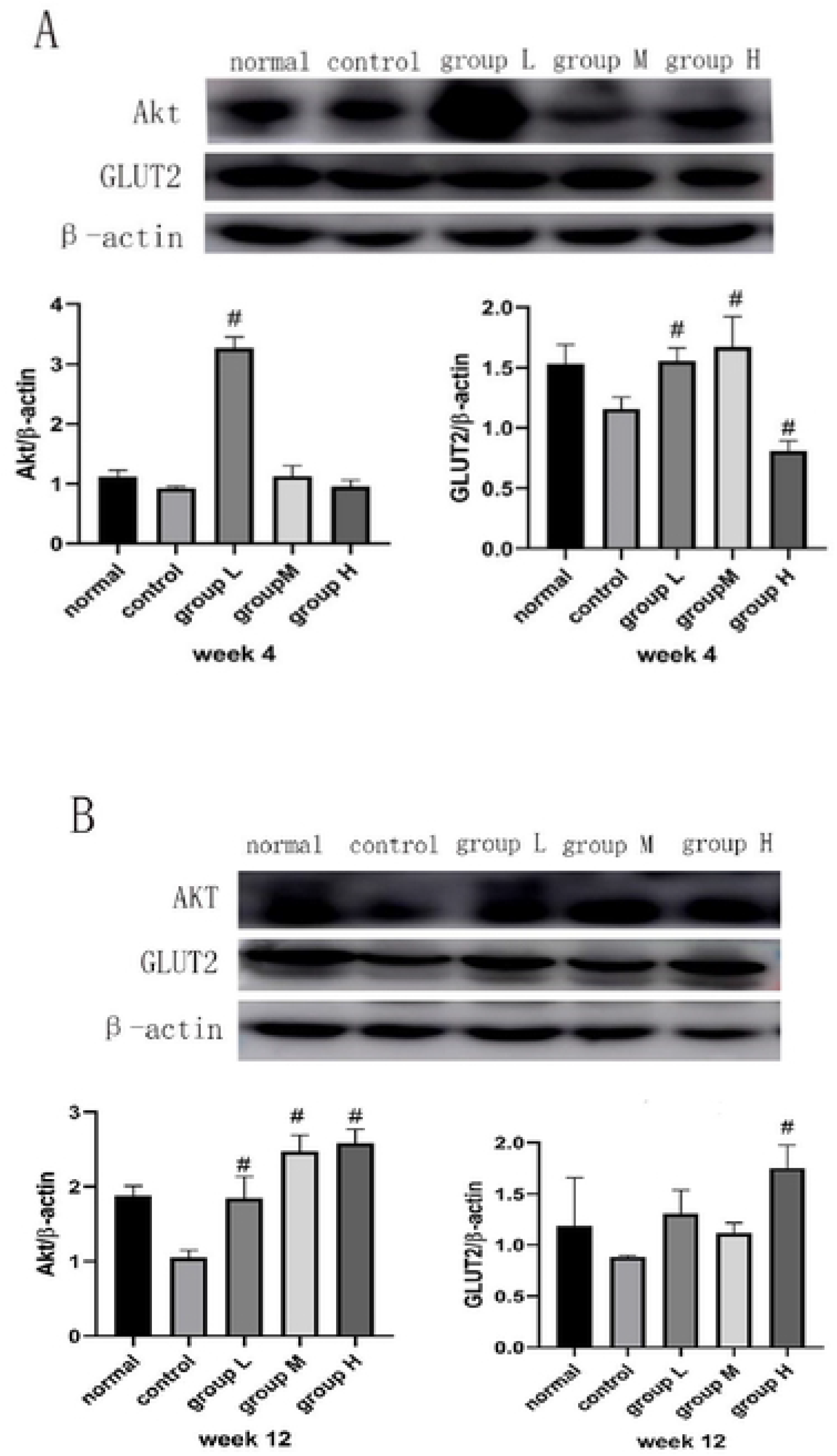

### Changes in GLUT2 and AKT expression levels in rat liver tissues

As described in (Fig 3), changes in the expression of AKT and GLUT2 in the liver tissue of diabetic rats were observed after treatment with different doses of 1,25(OH)_2_D_3_. AKT expression levels were significantly up-regulated by 4-week low-dose 1,25(OH)_2_D_3_ intervention. GLUT2 expression levels in all treatment groups were also up-regulated by the 4 weeks 1,25(OH)_2_D_3_ intervention. After 12 weeks of intervention, the expression of AKT in all intervention groups was obviously increased, and the level of GLUT2 in the high-dose intervention group was also significantly increased (Fig 3B).

**Fig 3.**
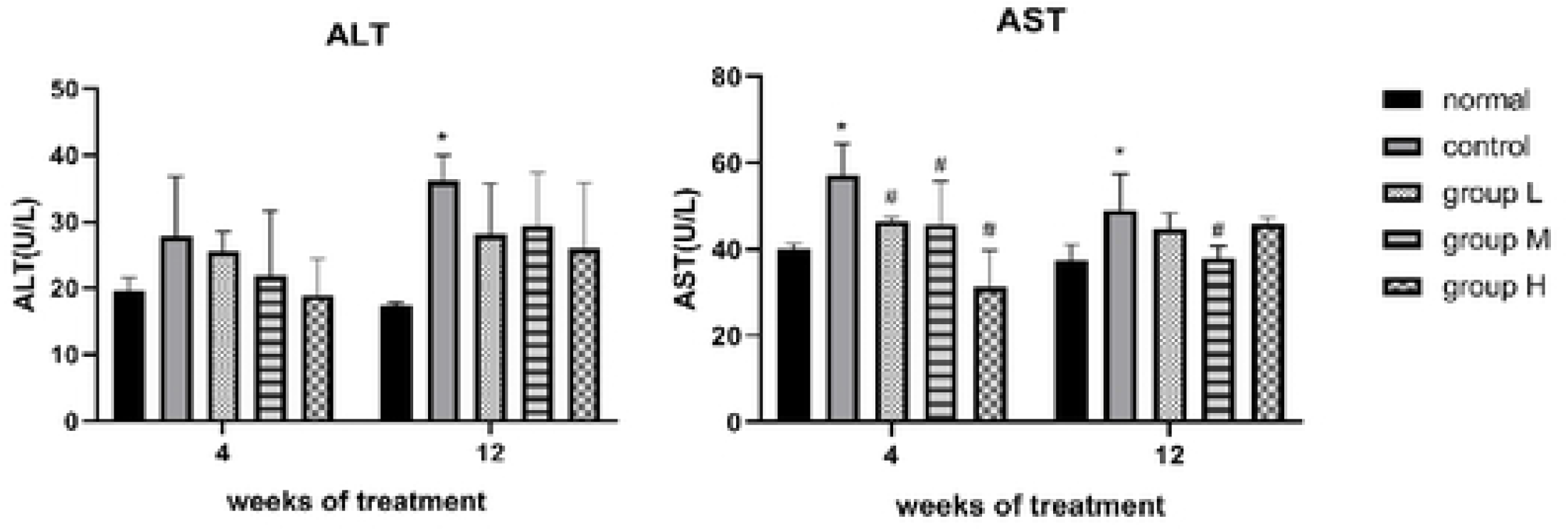

### Liver pathological changes of rats in each group

The liver tissue of normal rats was well structured, cells were fully shaped and neatly arranged. In the control group, the liver tissue of diabetic rats was disarranged with inflammation and vacuole. In the meantime, the pathological changes of liver was improved by 1,25(OH)_2_D_3_ intervention to a certain extent (Fig 4).

**Fig 4.**
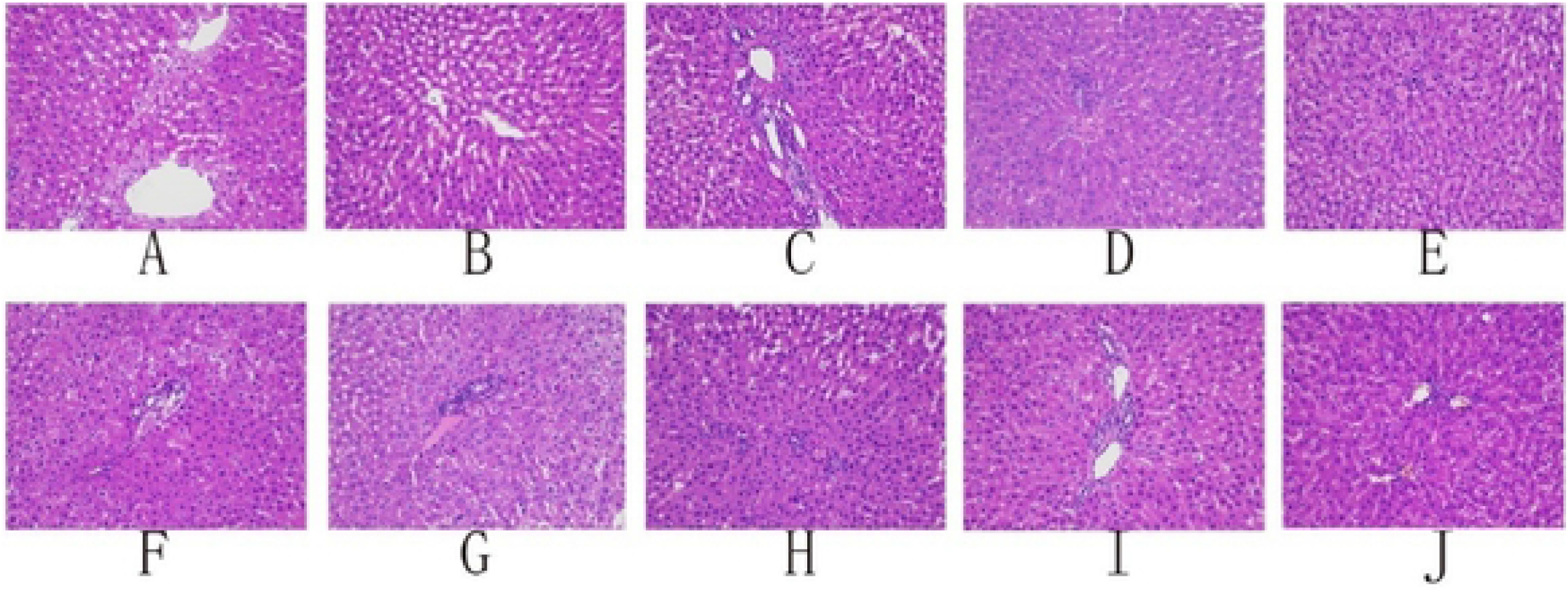
Effect of 1,25(OH)_2_D_3_ on the histological morphology of liver from rats. A,B,C,D and E were after 4 weeks intervention; F, G, H, I, J were after 12 weeks intervention. A and F were normal rats, B and G were control (type 2 diabetes) rats; C and H, type 2 diabetes rats treated with 1,25(OH)_2_D_3_of 0.075μg/Kg/d; D and I, type 2 diabetes rats treated with 1,25(OH)_2_D_3_ of 0.l50μg/Kg/d; E and J, type 2 diabetes rats treated with 1,25(OH)_2_D_3_ of 0.300μ g/Kg/d.

## Discussion

Diabetes is a group of chronic, systemic, and metabolic diseases characterized by hyperglycemia. Hyperglycemia is caused by insufficient insulin secretion and/or insulin resistance, which can lead to serious complications such as macroangiopathy, diabetic nephropathy, retinopathy, and diabetic foot [10]. Recently, diabetes and its complications have become global public health issues that require urgent solutions.

Vitamin D_3_ is derived from the food or synthesized in the skin under the influence of ultraviolet B light (UVB) through a photochemical reaction. Vitamin D_3_ obtained from the food and skin undergoes 25-hydroxylation in the human liver cells and is converted into 25(OH)D_3_, which is further converted into a more active 1,25(OH)_2_D_3_ in the kidney. 1,25(OH)_2_D_3_ combines with Vitamin D Receptor (VDR) widely present in various tissues and cells, thereby performing its biological functions [11]. Studies have shown that in addition to the classic calcium and phosphorus regulation, vitamin D is also involved in insulin resistance and immune regulation [12]. Epidemiological studies have revealed that patients with T2DM also have reduced serum vitamin D levels, and individuals with vitamin D deficiency have a higher risk of T2DM [13,14].

In this study, rats with STZ-induced type 2 diabetes were used to explore the mechanism of 1,25(OH)_2_D_3_ in improving their hepatic insulin resistance. A study by Wang et al. showed that lowering the level of insulin resistance could effectively improve lipid disorders and glucose tolerance in patients with type 2 diabetes [15]. In the present study, treatment with 1,25(OH)_2_D_3_ was found to reduce fasting blood glucose and increase the blood insulin levels in diabetic rats [16]. At the same time, it was observed that 1,25(OH)_2_D_3_ intervention could aid in treatment of lipid metabolism disorders, as indicated by the changes in serum contents of TC, TG, LDL-C, and HDL-C in this study.

The role of oxidative stress is evident in the pathogenesis of diabetes as well as in the occurrence and development of its complications [17]. Oxidative stress has long been implicated in inducing the apoptosis of islet cells, reducing the binding rate of insulin to its receptors and blocking insulin signal transduction, thereby leading to insulin resistance [18]. SOD is the main antioxidant enzyme that is widely present in the body and is responsible for removing the reactive oxygen species (ROS). MDA is the final product of lipid peroxidation, whose content can indirectly reflect the oxidative stress state of the body and the degree of cell injury caused by free radicals [19, 20]. In the present study, 1,25(OH)_2_D_3_ intervention was shown to increase the SOD activity in the liver tissue of diabetic rats, while reducing the content of MDA and improving the oxidative stress status of diabetic rats.

ALT and AST are biochemical markers of liver injury [21]. Previous studies have found that 1,25(OH)_2_D_3_ protects the rat liver from injury [22]. Intriguingly, our results showed no difference in the ALT levels among all groups; however, AST levels in the short-term (4-week) intervention groups were significantly lower than those in the control group. HE staining also showed that 1,25(OH)_2_D_3_ intervention could improve the pathological status of the liver in diabetic rats, indicating a protective role of 1,25(OH)_2_D_3_ in liver tissue lesions.

Glucose transporter 2 (GLUT2) is of great significance in improving glucose metabolism disorders in T2DM. GLUT2 is expressed in the liver, intestine, kidney, and pancreatic islet beta cells and in the central nervous system, neurons, astrocytes, and tanycytes. Physiological studies of genetically modified mice have revealed a role for GLUT2 in several regulatory mechanisms. In pancreatic beta cells, GLUT2 is required for glucose-stimulated insulin secretion [23]. A study by Yonamine et al. found that increased expression of GLUT2 is helpful for blood glucose control in diabetic rats [24]. AKT, a downstream effector of insulin and hepatic gluconeogenesis, functions as a key regulator of glucose uptake signaling [25, 26]. An earlier report by Carvalho et al. indicated that fat cells from T2DM patients tended to have lower expression of p-Akt and reduced translocation of plasma membrane Akt [27].

Moreover, a recent animal study found that the expression of GLUT2 in rats was dramatically downregulated by STZ treatment, whereas myrtenal administration significantly increased the expression of Akt in muscle and liver tissues while improving the diabetic status in rats [28]. Moreover, oxidative stress has been implicated as the main factor causing insulin resistance, and previous studies have shown that excessive ROS generation could induce T2DM by affecting insulin secretion, glucose utilization, and stimulating inflammation [29]. Thus, there is possibly a link between oxidative stress and the aberrant expression of p-Akt and GLUT2.

In the liver, Akt regulates hepatic glucose and lipid metabolism by affecting insulin signaling [30]. One of the mechanisms of hepatic insulin resistance is the phosphorylation of serine residues in the insulin receptor and Insulin Receptor Substrate-1 (IRS-1), thereby reducing the enzyme activity of the PI3K/AKT pathway ^31^. Treatment with 1,25(OH)_2_D_3_ significantly increased the expression of Akt and GLUT2 in rat liver, which may be due to the antioxidant effect of 1,25(OH)_2_D_3_ against lipid peroxidation.

Hepatic HE staining showed that after 1,25(OH)_2_D_3_ intervention, the degree of liver abnormalities in diabetic rats was reduced, probably due to the antioxidant function of 1,25(OH)_2_D_3_ that enhances the expression of Akt/GLUT2 protein in the liver and alleviates glucose and lipid metabolism disorders.

## Conclusion

In summary, our study demonstrated that treatment with 1,25(OH)_2_D_3_ increased the antioxidant capacity of diabetic rats by enhancing the SOD activity. It also promoted the expression of GLUT2 and AKT in the liver tissue, inhibited the decomposition of hepatic glycogen, ameliorated liver injury, improved insulin resistance, and alleviated the disorder of glucose and lipid metabolism in diabetic rats. Taken together, our results indicate that 1, 25(OH)_2_D_3_ might be a potential therapeutic agent for insulin resistance in diabetic liver.

## References

1. Hameed I, Masoodi SR, Mir SA, Nabi M, Ghazanfar K, Ganai BA. Type 2 diabetes mellitus: From a metabolic disorder to an inflammatory condition. World J Diabetes. 2015; 6(4): 598. doi:10.4239/wjd.v6.i4.598.

2. Rehman K, Akash MSH. Mechanism of Generation of Oxidative Stress and Pathophysiology of Type 2 Diabetes Mellitus: How Are They Interlinked? J Cell Biochem. 2017;118(11):3577–3585. doi:10.1002/jcb.26097.

3. Zhang JJ, Meng X, Li Y, et al. Effects of melatonin on liver injuries and diseases. Int J Mol Sci. 2017;18(4):1–27. doi:10.3390/ijms18040673.

4. Zand H, Morshedzadeh N, Naghashian F. Signaling pathways linking inflammation to insulin resistance. Diabetes Metab Syndr Clin Res Rev. 2017;11:S307–S309. doi:10.1016/j.dsx.2017.03.006.

5. Foroozanfard F, Talebi M, Samimi M, et al. Foroozanfard2017.

6. Manson JAE, Bassuk SS, Lee IM, et al. The VITamin D and OmegA-3 TriaL (VITAL): Rationale and design of a large randomized controlled trial of vitamin D and marine omega-3 fatty acid supplements for the primary prevention of cancer and cardiovascular disease. Contemp Clin Trials. 2012;33(1):159–171. doi:10.1016/j.cct.2011.09.009.

7. Mai S, Walker GE, Vietti R, et al. Acute vitamin D3 supplementation in severe obesity: Evaluation of multimeric adiponectin. Nutrients. 2017;9(5):1–14. doi:10.3390/nu9050459.

8. Komolmit P, Kimtrakool S, Suksawatamnuay S, et al. Vitamin D supplementation improves serum markers associated with hepatic fibrogenesis in chronic hepatitis C patients: A randomized, double-blind, placebo-controlled study. Sci Rep. 2017;7(1):1–11. doi:10.1038/s41598-017-09512-7.

9. Chandirasegaran G, Elanchezhiyan C, Ghosh K. Effects of Berberine chloride on the liver of streptozotocin-induced diabetes in albino Wistar rats. Biomed Pharmacother. 2018; 99 (September 2017): 227–236. doi:10.1016/j.biopha.2018.01.007.

10. Zheng Y, Ley SH, Hu FB. Global aetiology and epidemiology of type 2 diabetes mellitus and its complications. Nat Rev Endocrinol. 2018;14(2):88–98. doi:10.1038/nrendo.2017.151.

11. Borges MC, Martini LA, Rogero MM. Current perspectives on vitamin D, immune system, and chronic diseases. Nutrition. 2011;27(4):399–404. doi:10.1016/j.nut.2010.07.022.

12. Wacker M, Holiack MF. Vitamin D-effects on skeletal and extraskeletal health and the need for supplementation. Nutrients. 2013;5(1):111–148. doi:10.3390/nu5010111.

13. Dhas Y, Banerjee J, Damle G, Mishra N. Clinica Chimica Acta Association of vitamin D deficiency with insulin resistance in middle-aged type 2 diabetics. Clin Chim Acta. 2019;492(February):95–101. doi:10.1016/j.cca.2019.02.014.

14. Lim S, Kim MJ, Choi SH, et al. Association of vitamin D deficiency with incidence of type 2 diabetes in. Published online 2013:524–530. doi:10.3945/ajcn.112.048496.524.

15. Wang X, Xian T, Jia X, et al. A cross-sectional study on the associations of insulin resistance with sex hormone, abnormal lipid metabolism in T2DM and IGT patients. Med (United States). 2017;96(26). doi:10.1097/MD.0000000000007378.

16. Amin SN, Hussein UK, Yassa HD, Hassan SS, Rashed LA. Synergistic actions of vitamin D and metformin on skeletal muscles and insulin resistance of type 2 diabetic rats. J Cell Physiol. 2018;233(8):5768–5779. doi:10.1002/jcp.26300.

17. Rochette L, Zeller M, Cottin Y, Vergely C. Diabetes, oxidative stress and therapeutic strategies. Biochim Biophys Acta - Gen Subj. 2014;1840(9):2709–2729. doi:10.1016/j.bbagen.2014.05.017.

18. Droge W. Free radicals in the physiological control of cell function. Physiol Rev. 2002;82(1):47–95.

19. Fukai T, Ushio-Fukai M. Superoxide dismutases: Role in redox signaling, vascular function, and diseases. Antioxidants Redox Signal. 2011;15(6):1583–1606. doi:10.1089/ars.2011.3999.

20. Slatter DA, Bolton CH, Bailey AJ. The importance of lipid-derived malondialdehyde in diabetes mellitus. Diabetologia. 2000;43(5):550–557. doi:10.1007/s001250051342.

21. Otto-Ślusarczyk D, Graboń W, Mielczarek-Puta M. Aminotransferaza asparaginianowa--kluczowy enzym w metabolizmie ogólnoustrojowym człowieka. Postepy Hig Med Dosw (Online). 2016;70:219–230. doi:10.5604/17322693.1197373.

22. Özerkan D, Özsoy N, Akbulut KG, Güney, Öztürk G. The protective effect of vitamin D against carbon tetrachloride damage to the rat liver. Biotech Histochem. 2017;92(7):513–523. doi:10.1080/10520295.2017.1361549.

23. Thorens B. GLUT2, glucose sensing and glucose homeostasis. Diabetologia. 2015;58(2):221–232. doi:10.1007/s00125-014-3451-1.

24. Yonamine CY, Pinheiro-Machado E, Michalani ML, et al. Resveratrol Improves Glycemic Control in Type 2 Diabetic Obese Mice by Regulating Glucose Transporter Expression in Skeletal Muscle and Liver. Molecules. 2017;22(7). doi:10.3390/molecules22071180.

25. Kohn AD, Summers SA, Birnbaum MJ, Roth RA. Expression of a constitutively active Akt Ser/Thr kinase in 3T3-L1 adipocytes stimulates glucose uptake and glucose transporter 4 translocation. J Biol Chem. 1996;271(49):31372–31378. doi:10.1074/jbc.271.49.31372.

26. Leibiger B, Moede T, Uhles S, et al. Insulin-feedback via PI3K-C2α activated PKBα/Akt1 is required for glucose-stimulated insulin secretion. FASEB J. 2010;24(6):1824–1837. doi:10.1096/fj.09-148072.

27. Carvalho E, Eliasson B, Wesslau C, Smith U. Impaired phosphorylation and insulin-stimulated translocation to the plasma membrane of protein kinase B/Akt in adipocytes from type II diabetic subjects. Diabetologia. 2000;43(9):1107–1115. doi:10.1007/s001250051501.

28. Rathinam A, Pari L. Myrtenal ameliorates hyperglycemia by enhancing GLUT2 through Akt in the skeletal muscle and liver of diabetic rats. Chem Biol Interact. 2016;256:161–166. doi:10.1016/j.cbi.2016.07.009.

29. Biobaku F, Ghanim H, Batra M, Dandona P. Macronutrient-Mediated Inflammation and Oxidative Stress: Relevance to Insulin Resistance, Obesity, and Atherogenesis. J Clin Endocrinol Metab. 2019;104(12):6118–6128. doi:10.1210/jc.2018-01833.

30. Cai CX, Buddha H, Castelino-Prabhu S, et al. Activation of Insulin-PI3K/Akt-p70S6K Pathway in Hepatic Stellate Cells Contributes to Fibrosis in Nonalcoholic Steatohepatitis. Dig Dis Sci. 2017;62(4):968–978. doi:10.1007/s10620-017-4470-9.

31. Nandipati KC, Subramanian S, Agrawal DK. Protein kinases: mechanisms and downstream targets in inflammation-mediated obesity and insulin resistance. Mol Cell Biochem. 2017;426(1-2):27–45. doi:10.1007/s11010-016-2878-8.

